# Biosurfactant stabilized nanoemulsions as multifunctional magnetically targeted delivery vehicles

**DOI:** 10.1101/2023.11.26.568750

**Authors:** Russell J. Wilson, Yun Liu, Guangze Yang, Yuan Gao, Chun-Xia Zhao

## Abstract

Nanoemulsions have been widely used for pharmaceutic applications. However, there remains a significant challenge to functionalize available pharmaceutical surfactants with targeting or reporter moieties to generate next-generation drug delivery systems. Herein, a designed biosurfactant platform technology, based on our library of α-helical peptide AM1 derivatives, was used to prepare multifunctional magnetic nanoemulsion drug delivery systems. Key factors such as electrostatic and steric stabilization of the nanoemulsions were determined using this peptide library. Stabilization of the nanoemulsion was achieved by controlled and tunable loading of PEG-functionalized biosurfactant at the oil-water interface. A model drug and iron oxide nanoparticles were incorporated into the oil core to prepare multifunctional nanoemulsions. *In vitro* cell uptake experiments using an external magnetic field demonstrated controlled rapid uptake of iron oxide loaded nanoemulsions by SKOV3 cancer and RAW 264.7 macrophage cells. SKOV3 cells demonstrated a slower rate of uptake under a magnetic field when compared to RAW 264.7 cells. This work demonstrates a highly adaptable hierarchical nanoemulsion system that can provide a platform to develop effective multifunctional nanomedicines and tools for biomedical insights.

**Graphical Abstract:** 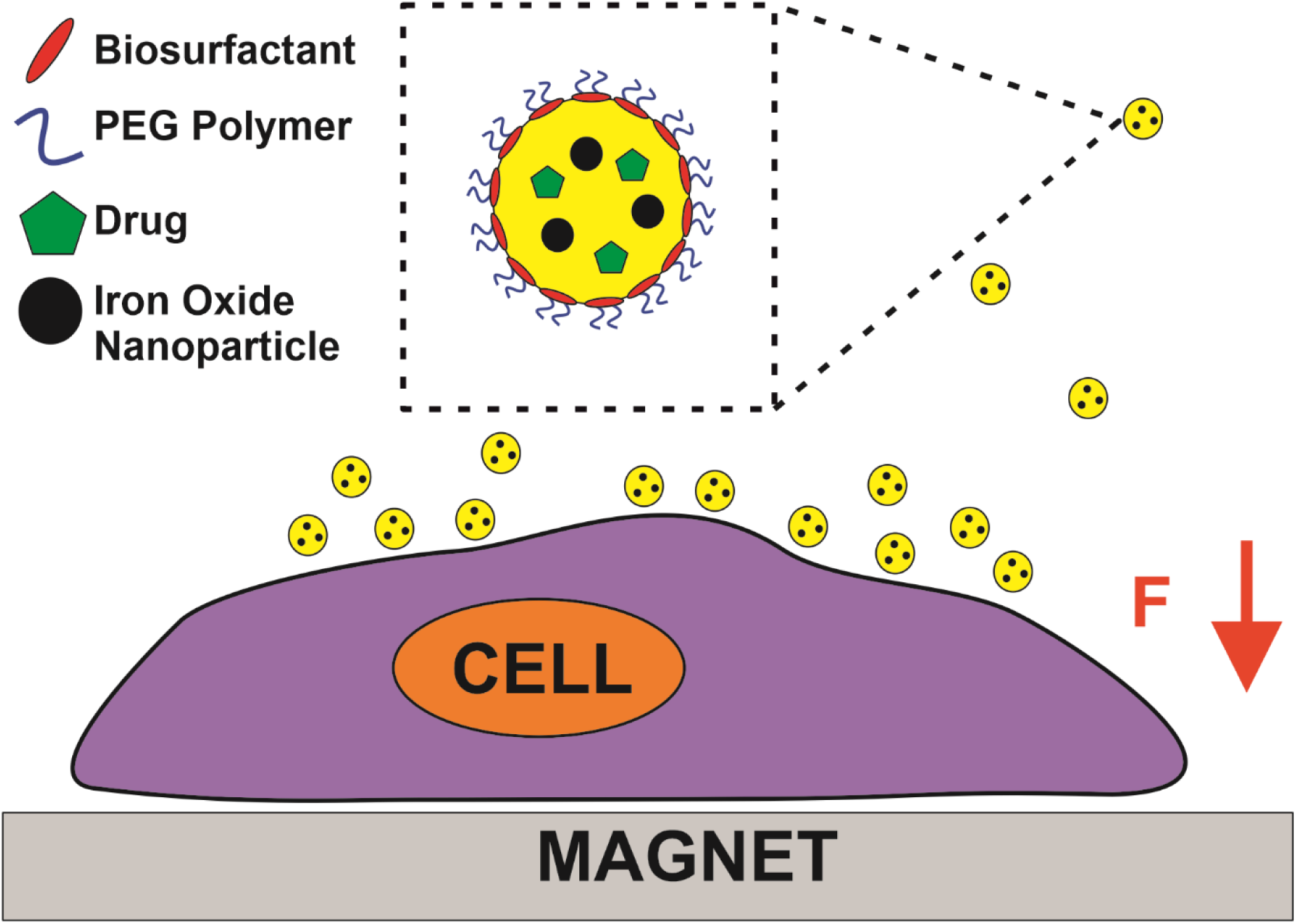

## Introduction

Nanoparticle based drug delivery vehicles have attracted great interest for cancer therapy in recent decades as they improve tumor accumulation, reduce off target effects and prolong circulation of poorly soluble drugs.[1] Spurred by greater understanding of the heterogeneity of cancers, tumors and biological barriers within and between patients, nanoparticle development has recently shifted towards personalized medicine.[2,3] From this a wide range of nanoparticles with multiple functionalities to better target, diagnose and treat disease have been developed.[4–6] However, many of the current systems lack flexibility in design for facile preparation combining features to efficiently and tunably load different drugs and functionalize the system with targeting, therapeutic and/or diagnostic moieties. These design factors must be readily and rapidly tailored to individual patient needs and are currently impeding the development pipeline of personalized nanomedicines. Thus, there is clearly a great need for high value, adaptable and facile nanoparticle platform technologies.

Protein based platform technologies are especially attractive for personalized nanomedicine as they are biocompatible, can self-assemble into complex nanostructures and have a plethora of strategies for functionalization. The highly specific biological functions of proteins can be exploited to form nanoparticles with unique physicochemical properties and therapeutic action.[7] This makes them distinct from the polymer, lipid and inorganic nanoparticle systems routinely investigated for drug delivery applications. Unfortunately to date protein based nanomedicines for drug delivery have seen limited clinical translation.[8] Abraxane® is the only currently in use FDA approved protein nanoparticle designed for drug delivery and is composed of albumin particles loaded with paclitaxel which significantly improves the safety and efficacy of breast cancer chemotherapy.[9] Due to the success of Abraxane® many of the current protein nanomedicines for drug delivery under preclinical investigation contain albumin, ferritin or transferrin proteins that are found in varying levels in the blood.[10] Using these proteins a range of protein nanocarriers have been developed to carry drug and/or diagnostic/therapeutic particles.[11,12] For example, Zhou et al. prepared a hierarchical nanocluster of albumin coated Gd^3+^ and CuS which was then encapsulated in ethylenediamine/phenylboronic acid functionalized dextran.[13] Using this system they demonstrated that extracellular ATP at the tumor site act triggered the release of the encapsulated material to provide contrast for magnetic resonance imaging and a sensitizer for photothermal therapy. Other protein based systems have utilized fibroin, lipoprotein and gelatin to delivery cargo to the cancer site.[14–16] Still there remains a significant challenge in the facile preparation of multifunctional protein nanomedicines that limits their clinical translation especially when integrating different organic and inorganic components with controlled and tunable loading.

Herein, we aim to investigate the key factors to develop a protein based multifunctional drug delivery vehicle using biosurfactants. To achieve this, we use a specially designed α-helical peptide sequence with surface active properties as a platform to identify key factors for protein stabilized nanoemulsions.[17] Using our library of biosurfactants with different chemical properties we develop series of multifunctional nanoemulsions containing iron oxide nanoparticles and test their stability as iron oxide nanoparticles can have surface active properties. Using this approach, multifunctional nanoemulsions were prepared by loading the oil core with hydrophobic iron oxide nanoparticles and a model drug. Overall, this protein based nanoemulsion platform offers new strategies for external and internal functionalization for nanomedicines with higher therapeutic efficacy and patient safety.

## Materials & Methods

### Materials

Miglyol-812 was received from Cremer Oleo (Hamburg, Germany) and purified by passing through heat-activated silica gel (Sigma-Aldrich, Castle Hill, Australia) prior to use. *N*-hydroxysuccinamide ester (NHS) functionalized polyethylene glycol (PEG-NHS) (average M_W_ = 5000 kDa, PDI <1.08, purity >95 %) was purchased from Nanocs (New York, USA). Paclitaxel (˃99.5%) was purchased from LC Laboratories (Woburn, MA). The synthetic peptides AM1 (theoretical M_w_ = 2472 g·mol^-1^) and SurSi (theoretical M_w_ = 3632 g·mol^-1^) were designed in our lab and synthesized by GenScript® (Nanjing, China) to a minimum 95% purity. Gibco™ Dulbecco’s Modified Eagle Medium (DMEM), Gibco™ Trypsin-EDTA 0.5% solution, Gibco™ fetal bovine serum (FBS) and DiI fluorescent dye were purchased from Thermofisher (MA, USA). Roche Cell Proliferation WST-1 reagent was purchased from Sigma-Aldrich (MI, USA). Ultrapure water having a resistivity of >18.2 MΩ.cm was produced from a Milli-Q water purification system (Merck Millipore, Bayswater, Australia). Isopropyl β-D-1-thiogalactopyranoside (IPTG) was purchased from AMRESCO® (Solon, US). All other reagents were purchased from Sigma-Aldrich and were of analytical grade.

### DAMP4 Bacterial Expression

Protein surfactant DAMP4 sequence was cloned into a pET48b vector and expressed in E. coli strain BL21 (DE3) (Protein Expression Facility, The University of Queensland, Australia). The bacteria from glycerol stock stored at -80°C was streaked onto LB agar plates (15 g·L^-1^ agar, 10 g·L^-1^ peptone, 5 g·L^-1^ yeast extract, 10 g·L^-1^ NaCl, 50 mg·L^-1^ Kanamycin) and grown overnight at 37 °C. A single colony was chosen and used to inoculate 10 mL of 2YT media (16 g·L^-1^ peptone, 10 g·L^-1^ yeast extract, 5 g·L^-1^ NaCl, 50 mg·L^-1^ Kanamycin) and grown overnight in an incubator at 180 rpm and 37 °C. 800 µL of starter culture was used to inoculate 800 mL of 2YT media and grown for 4 h in an incubator at 180 rpm and 37 °C. The cultures were then induced using 1 mM (IPTG) and further incubated for 4 h in an incubator at 180 rpm and 37 °C. The cells were collected by centrifugation (Avanti® JXN-30) at 4000 xg for 20 min at 4°C and cells pellets stored at -20°C until required.

### DAMP4 Purification

DAMP4 was purified with three sequential chromatographic methods namely: ion-metal affinity (IMAC), ion exchange (IEX) and reverse-phase high performance chromatography (RP-HPLC). The *E. coli* bacterial pellet stored at -20°C were resuspended in 40 mL of lysis buffer (50 mM NaCl, 25 mM Na_2_HPO_4_, 2 mM MgCl_2_, 0.5% w/v TritonX-100, pH 7.5) and their cell walls disrupted by ultrasonication (Branson Ultrasonics Corporation, Connecticut, USA) at power level 5 for four 45 second cycles with a 1 min rest on ice between each cycle. The cell lysate was centrifuged (Avanti® JXN-30) at 40,000 xg for 20 min at 4°C and the supernatant was filtered through a 0.45 µm Millex® syringe filter (Merck, Darmstadt, Germany). Clarified lysate was loaded onto a nickel (II) 5 mL HisTrap packed with Sepharose® 6 fast flow agarose beads (GE Healthcare Biosciences, NSW, Australia) using an ÄKTApure™ system (GE Healthcare Biosciences). The column was preequilibrated with 5 column volumes of Buffer A (50 mM NaCl, 25 mM NaH_2_PO_4_, pH 7.5) and loosely bound protein was eluted at 6% Buffer B (500 mM imidazole, 50 mM NaCl, 25 mM NaH_2_PO_4_, pH 7.5) and DAMP4 was eluted at 80% Buffer B. Elution fractions were combined, heated to 60°C to remove aggregates and further purified on two coupled IEX columns 1 mL HiTrap Q FF and 1 mL HiTrap SP FF (GE Healthcare Life Sciences, NSW, Australia) by collecting the flow-through. The flow-through was purified on a Jupiter C5 10 μm 300 Å 250 mm × 10 mm column (Phenomenex, NSW, Australia) using Buffer A (99.9% ultrapure water, 0.1% trifluoroacetic acid) and Buffer B (90% acetonitrile, 9.9% ultrapure water, 0.1% trifluoroacetic acid). The isolated DAMP4 was quantified using RP-HPLC with an established standard curve, lyophilized and stored at -80°C.

### DAMP4 PEGylation

Lyophilized DAMP4 was dissolved in HEPES buffer (25 mM, pH 7) and added to PEG-NHS at a molar ratio of 1:40. The products were left to stir at room temperature for 4 h and then lyophilized and stored at -80°C. To confirm PEGylation the product and reactants were analyzed by SDS-PAGE using Mini-PROTEAN® TGX™ Precast protein gels (Bio-Rad Laboratories Inc., CA, USA).

### AM1 Emulsion Preparation

Lyophilized AM1 and zinc chloride were made to a final concentration of 400 µM and 800 µM, respectively, in HEPES buffer (25 mM, pH 7). To make 1 mL nanoemulsion, 980 µL of AM1+ZnCl_2_ solution was added to 20 µL of Miglyol-812. For dye loaded experiments DiI was dissolved in Miglyol-812 oil at a concentration of 0.5 mg·mL-1. For iron oxide loaded nanoemulsions, prior to homogenization the Miglyol-812 was mixed with 20 µL 5 mg·mL^-1^ oleic acid coated iron oxide nanoparticles in toluene and heated to 110°C to remove toluene. The mixture was homogenized by ultrasonication (Branson Ultrasonics Corporation, Connecticut, USA) at power level 2 for five 30 second cycles with a 60 second rest on ice between each cycle. The nanoemulsions were stored at 4°C in low light conditions until required.

### SurSi Emulsion Preparation

Lyophilized SurSi and ZnCl_2_ were made to a final concentration of 400 µM and 800 µM, respectively, in HEPES buffer (25 mM, pH 7.5). To make 3 mL nanoemulsion, 2940 µL of SurSi+ZnCl_2_ solution was added to 60 µL of Miglyol-812. For iron oxide loaded nanoemulsions, prior to homogenization the Miglyol-812 was mixed with 20 µL 5 mg·mL^-1^ oleic acid coated iron oxide nanoparticles in toluene and heated to 110°C to remove toluene. The mixture was pre-homogenized using an overhead rotor stator homogenizer at power level 6 (Ystral, Ballrechten-Dottingen, Germany) for 60 seconds. The mixture was further homogenized by ultrasonication (Branson Ultrasonics Corporation, Connecticut, USA) at power level 2 for five 30 second cycles with a 60 second rest on ice between each cycle. The nanoemulsions were stored at 4°C in low light conditions until required.

### AM1 DAMP4 Nanoemulsion Preparation

An AM1 nanoemulsion was prepared as previously described. Lyophilized DAMP4 was dissolved in ultrapure water to a final concentration of 400 µM and a separate solution to a final concentration of 40 µM DAMP4 by a 1 in 10 dilution. A DAMP4_20_-AM1 nanoemulsion was prepared by adding AM1 nanoemulsion to 40 µM DAMP4 at a 1:1 ratio and mixed by inversion for 60 seconds. A DAMP4_20-200_-AM1 nanoemulsion was prepared by adding DAMP4_20_-AM1 nanoemulsion to 400 µM DAMP4 at a 1:1 ratio and mixed by inversion for 60 seconds. The nanoemulsions were stored at 4°C in low light conditions until required. DAMP4_20-200_-AM1 nanoemulsions are the working solution and referred to as DAMP4 AM1 nanoemulsions in the manuscript.

### AM1 DAMP4-PEG Nanoemulsion Preparation

An AM1 nanoemulsion was prepared as previously described. Lyophilized DAMP4-PEG was dissolved in ultrapure water to a final concentration of 400 µM and a separate solution to a final concentration of 40 µM DAMP4-PEG by a 1 in 10 dilution. A DAMP4-PEG_20_-AM1 nanoemulsion was prepared by adding AM1 nanoemulsion to 40 µM DAMP4-PEG at a 1:1 ratio and mixed by inversion for 60 seconds. A DAMP4-PEG_20-200_-AM1 nanoemulsion was prepared by adding DAMP4-PEG_20_-AM1 nanoemulsion to 400 µM DAMP4-PEG at a 1:1 ratio and mixed by inversion for 60 seconds. The nanoemulsions were stored at 4°C in low light conditions until required. DAMP4-PEG_20-200_-AM1 nanoemulsions are the working solution and referred to as DAMP4-PEG AM1 nanoemulsions in the manuscript.

### Particle Size & Zeta-Potential Determination

Size and zeta-potential of nanoemulsions was determined by analysis with a Malvern Zetasizer Nano ZS (Malvern, Worcestershire, UK). Solutions were diluted to approximately 100 times stock nanoemulsion concentration and 1 mL of the diluted solution was added to a 4 mL polystyrene cuvette (Sarstedt, Nümbrecht, Germany) or a 1 mL folded capillary cell (Malvern, Worcestershire, UK) and analyzed at 0 and 24 h. For the FBS analysis 125 µL of fresh fetal bovine serum was added to 375 µL of nanoemulsion and mixed for 60 seconds by inversion. Then 12.5 µL of FBS nanoemulsion solution was diluted by addition of 1 mL of water and analyzed at 0 and 24 h. For the DMEM analysis 800 µL of DMEM supplemented with FBS and dual antibiotics was added to 200 µL of nanoemulsion and mixed for 60 seconds. Then 50 µL of DMEM nanoemulsion solution was diluted by addition of 1 mL of water and analyzed at 0 and 24 h.

### Dye Release

The dye release of DiI loaded DAMP-PEG nanoemulsions was assessed in stock solution at 4 °C at 0 and 24 h. At the time point 20 µL of sample was taken and replaced with an equivalent volume of HEPES buffer. The sample was diluted 20 times and analyzed using a plate reader (Tecan, Zürich, Switzerland) and excitation wavelength λ_ex_ = 550 nm and emission wavelength λ_em_ = 580 nm.

### Cell Culture

RAW 264.7, primary human mammary Fibroblast and SKOV3 cells were purchased from American type culture collection (ATCC). RAW 264.7, human mammary Fibroblast and SKOV3 cells were cultured in DMEM medium supplemented with 10% v/v FBS and 1% (v/v) penicillin streptomycin at 37 °C with a humidified 5% CO_2_ atmosphere.

### Iron Oxide Cytotoxicity Study

A day prior to the uptake experiment, RAW 264.7, Fibroblast or SKOV3 cells were added to a Costar® flat bottom 96 well plate (Corning, New York, United States) at a density of 1 × 10^4^ cells per well with a layer of sacrificial PBS added around the cells to prevent evaporation and incubated at 37 °C with 5% v/v CO_2_. For the cytotoxicity experiment, the medium was removed from each well and replaced with 10 µL of HEPES buffer (25 mM, pH 7), DAMP4-PEG_20-200_-AM1 nanoemulsion or iron oxide loaded DAMP4-PEG_20-200_-AM1 nanoemulsion mixed with 190 µL of culture medium. The cells were incubated for 0.5, 2 or 4 h and for magnetically driven uptake the 96 well plate was placed on top of a magnetic plate (Ozbiosciences, Marseilles, France) for the duration. Once completed the medium was removed and replaced with 20 µL of WST-01 reagent mixed with 200 µL of culture medium and incubated the cells for 2 h. The cells were immediately analyzed using a plate reader (Tecan, Zürich, Switzerland) at 450 nm.

### *In Vitro* Cell Uptake

A day prior to the uptake experiment, RAW 264.7 or SKOV3 cells were added to a Costar® flat bottom 24 well plate (Corning, New York, United States) at a density of 5 × 10^5^ and 1 × 10^5^ cells per well respectively and incubated at 37 °C with 5% v/v CO_2_. For the uptake experiment, the medium was removed from each well and replaced with 1 mL of 50 µL of HEPES buffer (25 mM, pH 7), DiI loaded DAMP4-PEG_20-200_-AM1 nanoemulsion or iron oxide and DiI loaded DAMP4-PEG_20-200_-AM1 mixed with 950 µL of culture medium. The cells were incubated for 0.5, 2 or 4 h and for magnetically driven uptake the 24 well plate was placed on top of a magnetic plate (Ozbiosciences, Marseilles, France) for the duration. Once completed the medium was removed and the cells washed with PBS then dissociated with trypsin (0.25%). The cells were washed 3 times and resuspended in PBS. The cell uptake of nanoparticles were analyzed by flow cytometry using a CytoFLEX™ flow cytometer (Beckman Coulter, California, USA). For confocal imaging, cells were treated for 2 h in the presence or absence of a magnetic field. The cells were then washed twice with PBS, treated with a 4% formaldehyde solution for 15 min and then washed again with PBS. The cells were imaged on a Zeiss LSM 710 confocal laser scanning microscope (Carl Zeiss AG, Oberkochen, Germany).

### Data Analysis

Experiments were performed with at least three independent repetitions. Statistical analysis was conducted using GraphPad Prism 9 software. Two-tailed Student’s t-test were used to determine statistical significance and statistical differences were defined as *P < 0.05, **P < 0.01, ***P < 0.001 and ****P < 0.0001.

## Results & Discussion

### Biosurfactant Design

Protein biosurfactants are a versatile platform with high biocompatibility, low immunogenicity and special structure-function relationship that can be used to develop an array of nanoemulsion formulations. The design of amino acid sequences with specialized functions can be imparted via click chemistry. For example, introducing cysteine or lysine residues into the protein structure provide bioconjugation opportunities to form disulfide linkers, maleimide thioethers or *N*-hydroxysuccimide (NHS) amides. Fusion proteins can be designed by recombinant DNA and protein technology to express protein biosurfactants with unique biological functions such as integrated targeting peptides, enzymatic behavior and/or therapeutic monoclonal antibodies. This sets them apart from zwitterionic or non-ionic surfactants routinely used in clinical formulations. Pharmaceutical surfactants, such as lecithin, Tween and Cremophor EL, have limited functionality, regulatory limitations and/or require extensive chemical modification to add targeting/therapeutic moieties. Thus, designing protein based biosurfactants with favorable properties for drug delivery systems is highly attractive.

For drug delivery applications, nanoemulsions must have strong interfacial stabilization to withstand the complex biological environment of the blood and reach the intended target. An ideal protein biosurfactant must also provide stability during facile preparation of complex hierarchical materials. As such we considered the AM1 peptide (**Table S1**) (Ac-MKQLADS LHQLARQ VSRLEHA-CONH_2_) that conforms to an α-helical secondary structure as a platform to evaluate 4 different biosurfactant systems (**Figure 1a & 1b**). The designed sequence incorporates hydrophobic residues on one side and hydrophilic on the other that allows the peptide sit along an oil-water interface of a nanoemulsion droplet. The two specifically located histidine (H) residues in the peptide sequence provide an additional stabilizing force by forming an interconnected Zn^2+^ bridge network between individual peptides. This interfacial network provides a strong physical barrier to stabilize the droplets against coalescence (the merging of two or more droplets) and is not present for typical chemical surfactants. In addition, surface charge and electrostatic stabilization play two important roles in nanoparticle drug delivery systems improving the stability of the particle suspension by providing repulsive forces between particles and regulating uptake behavior.

**Figure 1.**
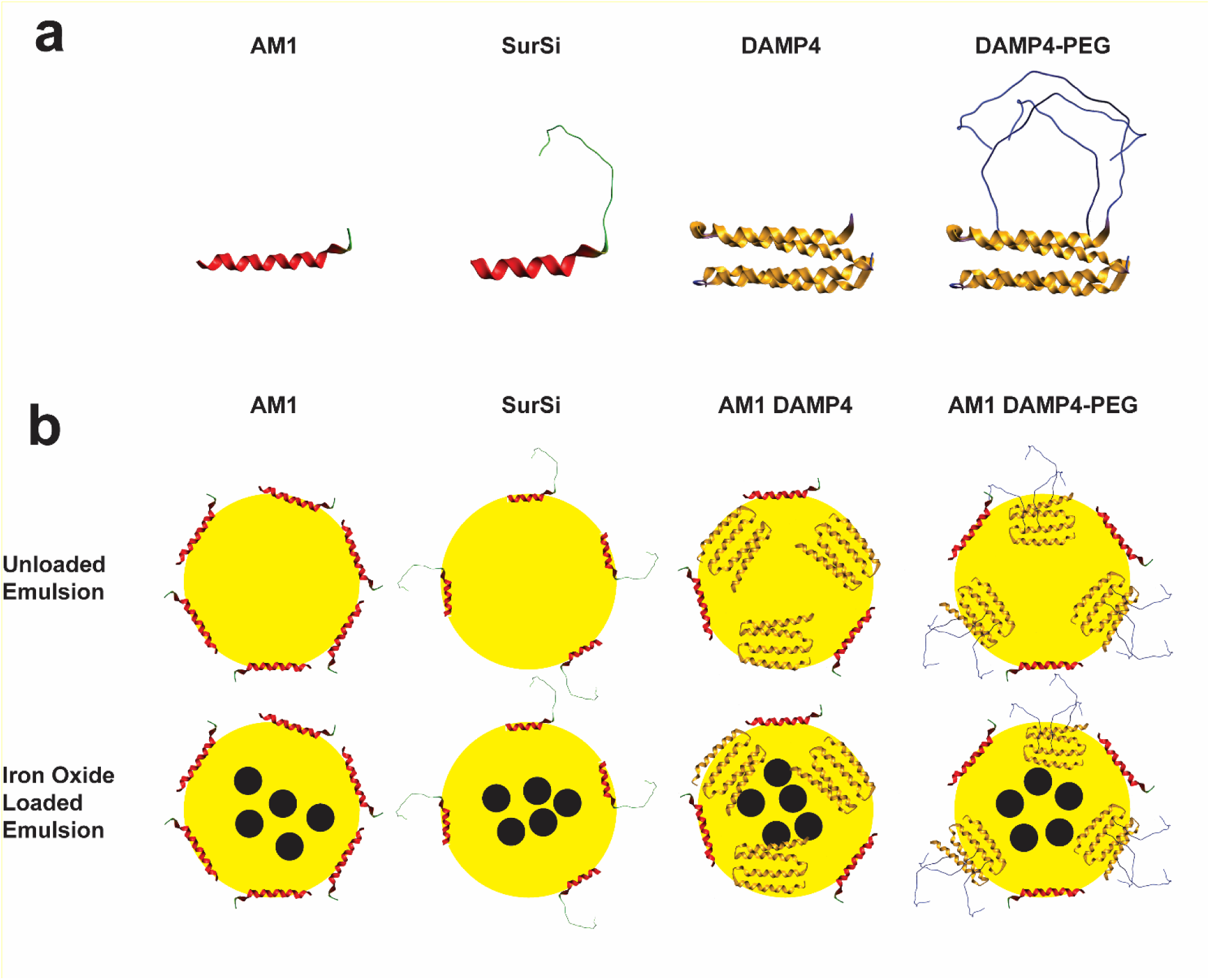
Illustration of biosurfactants and stabilized nanoemulsion systems. (a) Designed biosurfactants AM1, SurSi, DAMP4 and DAMP4-PEG. (b) Unloaded and iron oxide loaded nanoemulsions prepared using each biosurfactant.

To explore the influence of surface charge we used an AM1 peptide derivative called SurSi (Ac-MKQLAHS VSRLEHA RKKRKKRKKRKKGGGY-CONH_2_) that contains a random coil sequence rich in the positively charged amino acids arginine (R) and lysine (K).[18] The positively charged random coil sequence provides enhanced electrostatic repulsion between like-charged nanoemulsion droplets. Finally, an important translational consideration for biosurfactant design is their synthesis which is typically expensive and poorly scalable when using chemical methods. Designing biosurfactants for biosynthesis using an expression vector in living cells allows scalable production and specificity in function that is not readily achieved with chemical synthesis. To that end we used DAMP4 (MD(PSMKQLADS LHQLARQ VSRLEHAD)_4_) as a model biosurfactant protein that is prepared using a bacterial expression vector.[19] DAMP4 is composed of four AM1 peptides joined using a DPS (aspartic acid, proline, serine) linker and conforms to a 4-helix bundle structure. When exposed to an oil-water interface the 4-helix bundle structure of DAMP4 can unfold to expose the hydrophobic core and sit along the interface similar to AM1. It is important to note that a significant barrier to utilizing 4-helix bundle biosurfactants is that the slow kinetics of helix bundle unfolding directly impacts the stability of nanoemulsion droplets. The amine groups of the lysine residues in DAMP4 are also perfect locations for NHS esterification with additional functionality such as PEG polymer attachment. Attaching hydrophilic polymers such as PEG to a biosurfactant provide additional steric interfacial stabilization during nanoemulsion preparation and improves their efficacy and biodistribution during drug delivery.

### Multifunctional Nanoemulsion Formation & Stability for Biological Applications

An effective nanoemulsion drug delivery system must have good size and stability in biological media.[20] We prepared eight different nanoemulsion systems using the four peptides & proteins introduced previously and evaluated their physical properties in buffer, FBS or DMEM. For four of the nanoemulsion systems we preloaded oleic acid coated iron oxide nanoparticles with an average size of 20 nm into the oil phase. During the formation of the nanoemulsion systems droplet size is controlled by several factors namely surfactant adsorption kinetics, surfactant concentration and shearing energy. We maintained a consistent energy input and surfactant concentration (400 µM) during ultrasonication allowing us to investigate the influence of each surfactant. The surfactant concentration was chosen as we have previously found that at and beyond this concentration the relationship between droplet size and biosurfactant concentration is less significant.

SurSi and AM1 O/W nanoemulsions were formed by ultrasonication of a small volume of oil in buffer containing biosurfactant **(Figure 2a)**. For the DAMP4 and DAMP4-PEG nanoemulsions we developed a sequential addition method by adding DAMP4 derivatives to AM1 nanoemulsions **(Figure 2b)**. DAMP4-PEG was produced by reaction with 5 kDa N-hydroxysuccinimide ester functionalized poly(ethylene glycol) (NHS-PEG) and the conjugation confirmed by SDS-PAGE analysis **(Figure S1)**. DAMP4 derivatives alone form less monodisperse and larger nanoemulsions due to their slow unfolding kinetics at the interface. The strong surface activity of AM1 allows the formation of an monodisperse nanoemulsion template. To the template a dilute quantity of DAMP4 derivative is added to initially stabilize the nanoemulsion during the slow adsorption of DAMP4. Next we add a higher concentration of DAMP4 to significantly increase the transfer of DAMP4 to the interface and further improve the stability. This procedure can also be exploited to controllably add different ratios of modified biosurfactant to the interface for example DAMP4-Ab-PEG and DAMP4-PEG.

**Figure 2.**
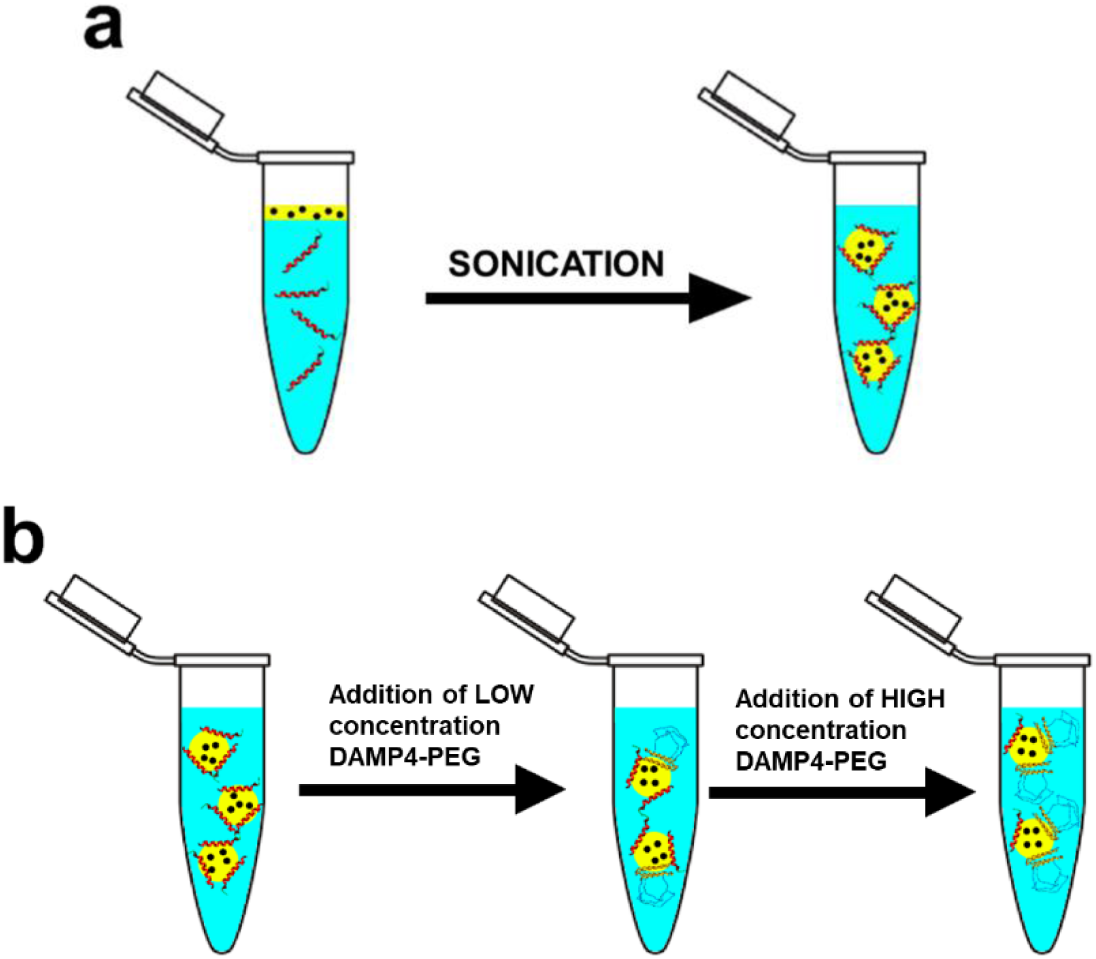
Formation of iron oxide loaded AM1 and AM1 DAMP4-PEG nanoemulsions. (a) Ultrasonication to form using AM1 and SurSi O/W nanoemulsion. (b) Sequential addition of DAMP4-PEG to AM1 nanoemulsion.

We first investigated the size and zeta potential of each nanoemulsion over 24 h in HEPES buffer **(Figure 3A & B)**. The AM1 and iron oxide loaded AM1 stabilized nanoemulsions had a size of 182 ± 2 and 197 ± 4 nm, respectively. They showed no significant change over the course of 24 h. However, we observed a significant difference between the zeta potentials of the unloaded and iron oxide loaded AM1 nanoemulsions being 50 ± 4 and 27 ± 3 mV, respectively. This reduction in zeta-potential is a result of the adsorption of the negatively charged iron oxide nanoparticle or oleic acid to the oil-water interface. This is significant as zeta-potentials above 40 lead to strongly electrostatically stable nanoemulsions, below 10 mV are prone to flocculation and between these are moderately stable. The SurSi and iron oxide loaded SurSi nanoemulsions had sizes of 202 ± 14 and 215 ± 2 nm, respectively, with no significant change over 24 h. We observed a significant increase in the zeta-potential of the iron oxide loaded SurSi nanoemulsion compared to the unloaded SurSi nanoemulsion being 42 ± 2 and 52 ± 3 mV, respectively. The zeta potential of iron oxide loaded SurSi nanoemulsions did not decrease as seen in the AM1 nanoemulsions potentially due to a combination of factors. The random coil of the Si module presenting the repeated RKK sequence may provide an electrostatic shield at the slipping plane. Also, the negative charged iron oxide nanoparticles may allow the adsorption of additional positive charged SurSi molecules to the surface. The degree of SurSi adsorption to an interface is controlled by strong electrostatic repulsion of nearby positively charged Si modules.

**Figure 3.**
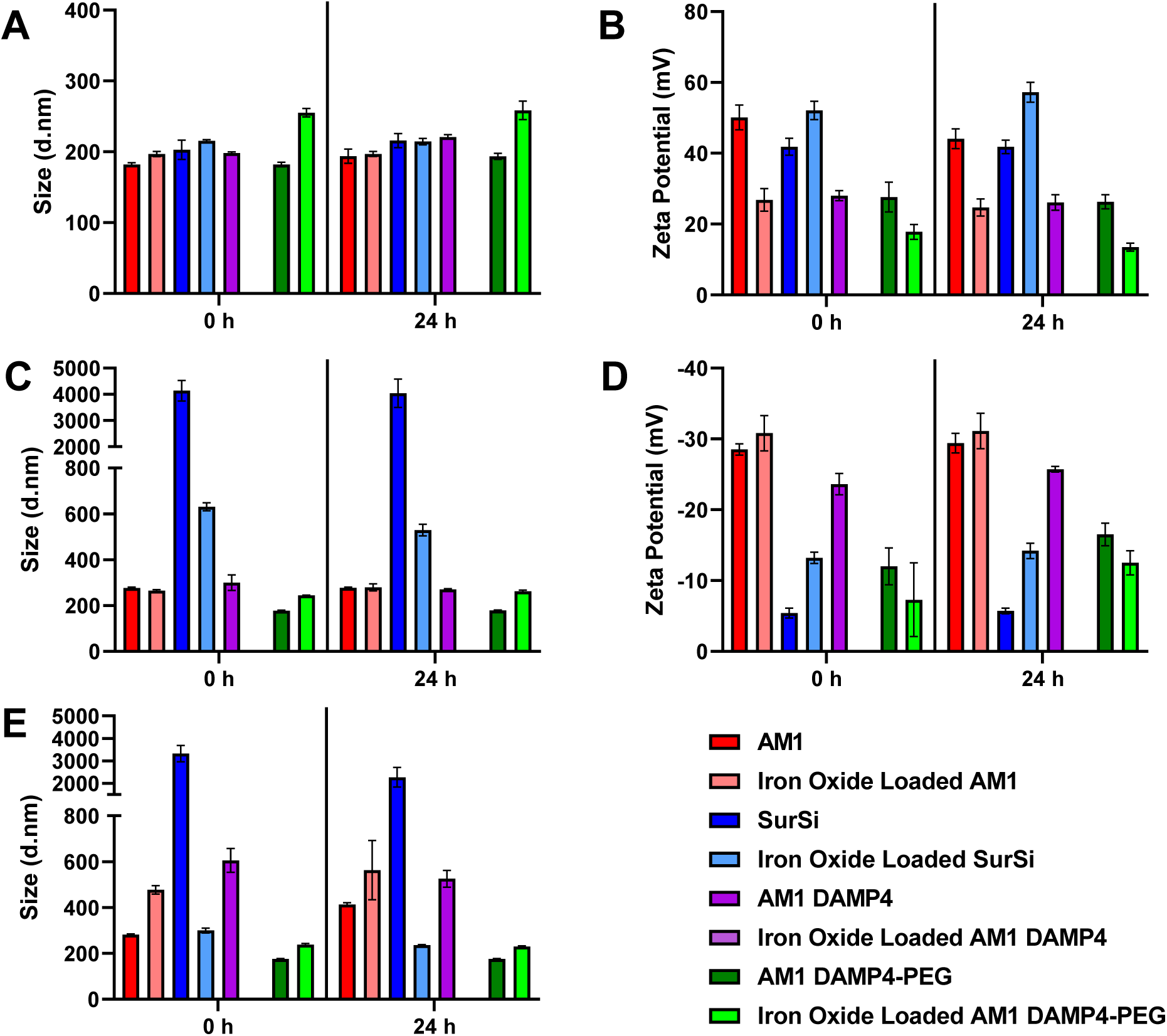
The size and zeta-potential data of biosurfactants stabilized nanoemulsions over 24 h (A) size in HEPES buffer (B) zeta-potential in HEPES buffer (C) size in 25% FBS (D) zeta-potential in 25% FBS (E) size in 80% DMEM supplemented with 10% FBS and 1% penicillin streptomycin. Note: Iron oxide loaded AM1 DAMP4 sample flocculated and is not shown. In all presented data the mean + s.d. from 3 separate replicates is shown.

For the AM1 DAMP4 nanoemulsion we observed a significant difference between the 0 and 24 h sample with sizes of 198 ± 2 and 221 ± 3 nm respectively. The zeta-potential of the AM1 DAMP4 was 28 ± 1 mV significantly lower than the pure AM1 nanoemulsions. This caused a small increase in size due to flocculation during the slow adsorption of the DAMP4 protein onto the surface of the nanoemulsion. This is because at pH 7 the DAMP4 protein has a negative charge from the additional aspartic acid residues in the four Asp-Pro-Ser linker sequences not present in AM1. The iron oxide loaded AM1 DAMP4 nanoemulsion experienced immediate flocculation into large aggregates and slowly creamed preventing assessment of the size and zeta-potential. This indicated the zeta-potential of the iron oxide loaded AM1 nanoemulsion decreased from 27 mV to near 0 mV upon the addition of the DAMP4 protein to the AM1 nanoemulsion. Further analysis of the iron oxide loaded AM1 DAMP4 nanoemulsion in biological media was not possible. The AM1 DAMP4-PEG nanoemulsion were significantly different with sizes 182 ± 3 and 194 ± 4 nm at 0 and 24 h, respectively. This difference was less than that observed for the unPEGylated DAMP4 protein as the PEG provides a steric barrier to flocculation and can screen the surface charge of the protein. The zeta-potential of the AM1 DAMP4-PEG nanoemulsion was 27 ± 4 mV, similar to that for the AM1 DAMP4 nanoemulsion. The iron oxide loaded AM1 DAMP4-PEG nanoemulsion was kinetically stable in contrast to the iron oxide loaded AM1 DAMP4 nanoemulsion. We measured a size of 255 ± 6 and 258 ± 13 nm at 0 and 24 h respectively, indicating that the PEG providing additional steric stability against flocculation. The zeta-potential of the iron oxide loaded AM1 DAMP4-PEG nanoemulsion was 18 ± 2 mV at 0 h and experienced a significant decrease to 14 ± 1 mV at 24 h. The zeta-potential of the iron oxide loaded AM1 DAMP4-PEG indicated poor electrostatic stabilization providing further evidence for why the unPEGylated protein induced flocculation.

Next we investigated the size and zeta-potential of each nanoemulsion system in 25% FBS to understand their stability under the influence of a protein corona **(Figure 3C & D)**. This is significant for nanomaterials used for drug delivery as the protein corona regulates physiological behavior. In addition, the adsorbed proteins can promote the formation of floc aggregates between the negatively charged proteins and positively charged nanoemulsions. The AM1 and iron oxide loaded AM1 nanoemulsions showed a significant increase in size to 276 ± 5 and 265 ± 4 nm, respectively, when dispersed in 25% FBS compared to pure buffer. The zeta potential of each nanoemulsion was -28 ± 1 and -31 ± 2 mV respectively showing the formation of a strongly negatively charged protein corona. There was no significant change in the size for either of the AM1 nanoemulsions over the course of 24 h due to the strong electrostatic repulsion. The SurSi nanoemulsion was significantly less stable in 25% FBS than the iron oxide loaded SurSi nanoemulsion with sizes of 4137 ± 393 and 630 ± 17 nm, respectively. The zeta potential of the SurSi nanoemulsion was -5.4 ± 1 mV and the iron oxide loaded SurSi nanoemulsion -13.2 ± 1 mV. These results demonstrate that the iron oxide loaded SurSi nanoemulsion was slightly stable due to moderate electrostatic repulsion, however, the SurSi nanoemulsion was poorly electrostatically stabilized and flocculated.

The AM1 DAMP4 nanoemulsion in 25% FBS was also significantly different to the AM1 DAMP4 in buffer increasing in size to 300 ± 34 nm. The AM1 DAMP4 nanoemulsion showed no significant difference over 24 h. The zeta potential of the AM1 DAMP4 nanoemulsion was -24 ± 2 mV indicating a moderate degree of electrostatic stabilization. The AM1 DAMP4-PEG nanoemulsion in FBS showed no significant difference compared to in HEPES buffer with a size of 178 ± 1 nm. The zeta potential of the AM1 DAMP4-PEG nanoemulsion was - 12 ± 3 mV and did not significantly change over 24 h. The iron oxide loaded AM1 DAMP4-PEG nanoemulsion showed a significant difference compared to HEPES buffer at 0 h with diameter of 244 ± 1 nm but no significant difference at 24 h with size of 262 ± 4 nm. The zeta potential of the iron oxide loaded AM1 DAMP4-PEG nanoemulsion was -7 ± 5 mV and did not significantly change over 24 h. The zeta potentials of the AM1 DAMP4-PEG and iron oxide loaded AM1 DAMP4-PEG nanoemulsion after 24 h of protein corona maturation showed a significant difference being -16.5 ± 2 and -12.5 ± 2 mV, respectively. The relatively low zeta-potentials did not significantly impact the stability of either of the AM1 DAMP4-PEG nanoemulsions as the steric stabilization of PEG provided an effective barrier to coalescence and flocculation.

Lastly, we observed the size of the nanoemulsion systems in 80% DMEM supplemented with 10% FBS and 1% dual antibiotic **(Figure 5)**. This allowed the evaluation of the stability in near isotonic conditions with a protein corona similar to that used in cell culture. The AM1 nanoemulsion increased from 281 ± 3 to 413 ± 6 nm and the iron oxide loaded nanoemulsion from 477 ± 18 to 563 ± 129 nm. The AM1 nanoemulsions showed a significant increase in size over time indicating some coalescence or flocculation in response to the high ionic strength media. High ionic strength can significantly alter the zeta potential of nanoparticles such that they can readily aggregate as the electrostatic repulsion is minimized. The SurSi nanoemulsion experienced less destabilization in 80% DMEM than that observed when in 25% FBS with a size of 3322 ± 363 nm at 0 h decreasing to 2266 ± 438 nm at 24 h. The iron oxide loaded SurSi nanoemulsion was significantly more stable in 80% DMEM than in 25% FBS with a size of 300 ± 11 nm at 0 h and decreasing to 236 ± 2 nm at 24 h.

**Figure 5.**
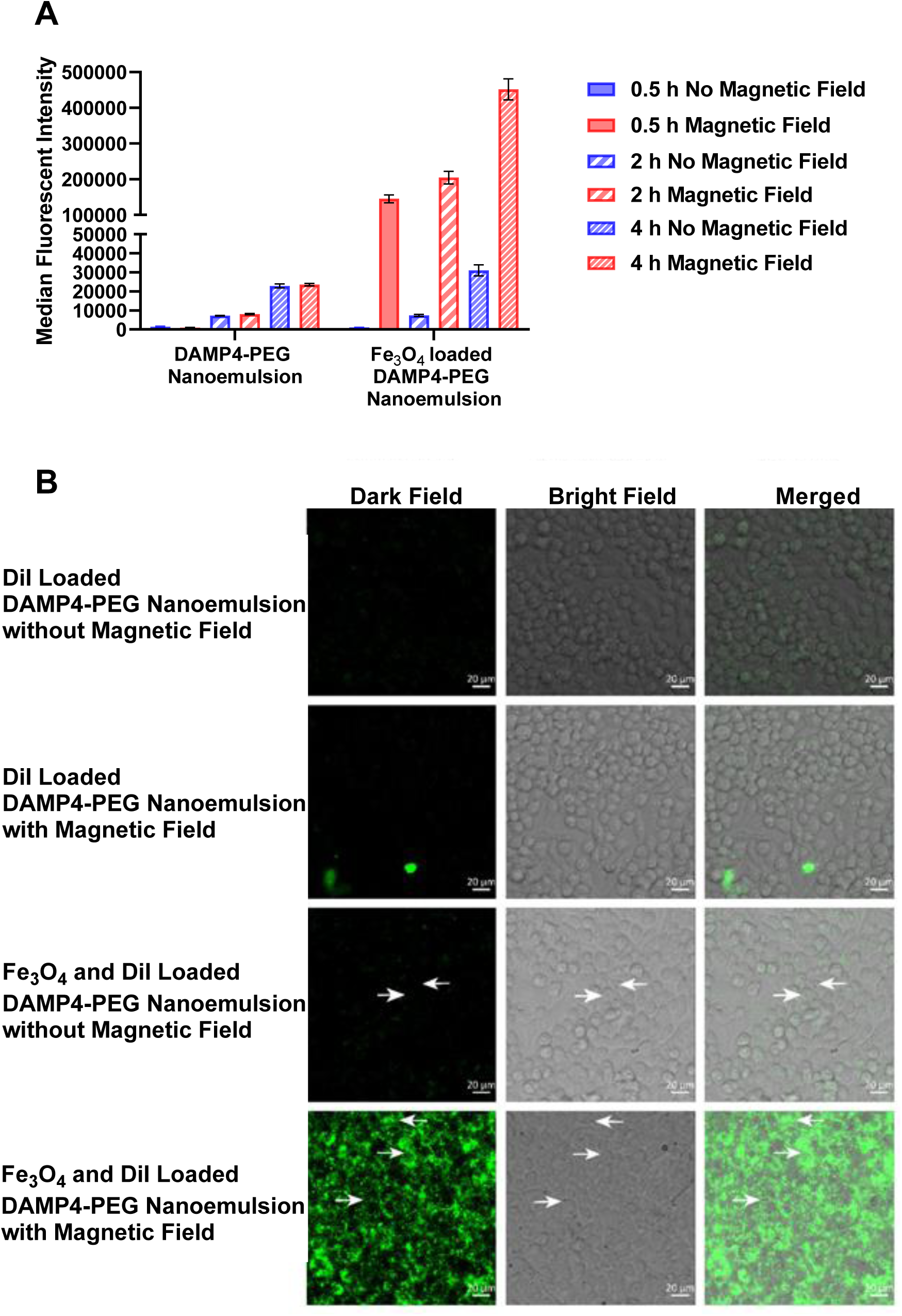
The cell uptake of DAMP4-PEG and iron oxide loaded DAMP4-PEG nanoemulsions by RAW 264.7 cells. **(A)** Flow cytometric data of the median fluorescent intensities at different time points in the presence and absence of a magnetic field. **(B)** Representative confocal images showing the cell uptake of DAMP4-PEG loaded nanoemulsions by RAW 264.7 cells in the presence and absence of a magnetic field after 2 h of incubation. Arrows indicate granular bodies. The mean + s.d. from 6 separate replicates is shown.

The AM1 DAMP4 nanoemulsion showed a significant increase in size to 605 ± 52 nm in 80% DMEM compared to in 25% FBS. This further demonstrated the significant impact of high ionic strength on the stability of the nanoemulsion. The AM1 DAMP4-PEG nanoemulsion showed no significant difference over time with a size of 175 ± 1 nm when dispersed in 80% DMEM. There was also no significant difference compared to in buffer or 25% FBS conditions. The iron oxide loaded AM1 DAMP4-PEG nanoemulsion showed no significant difference over time in 80% DMEM with a size of 239 ± 5 nm. Similarly, there was no significant difference between the sizes of the nanoemulsion when dispersed in FBS compared to DMEM. However, we observed a significant difference between the iron oxide loaded AM1 DAMP4-PEG nanoemulsion measured in buffer and DMEM at 0 h. This demonstrates the importance of the steric stabilization of PEG to prevent flocculation and regulate the formation of the protein corona.

Finally, as a core-shell material capable of loading hydrophobic cargo we needed to evaluate the capacity for the nanoemulsions to retain the cargo for extended periods of time. To do this we utilized the hydrophobic fluorescent dye DiI (λ_ex_ 550 nm, λ_em_ 580 nm) by dissolving it in the oil phase prior to emulsification. Over the course of 24 h the relative fluorescent intensity of the DAMP4-PEG nanoemulsion in stock solution decreased by 10% (**Figure S2**). Whereas, the relative fluorescent intensity of the iron oxide loaded DAMP4-PEG nanoemulsion in stock solution decreased by 22% over 24 h. It must also be noted that the total fluorescent intensity of the iron oxide loaded DAMP4-PEG nanoemulsion was 38% less than the DAMP4-nanoemulsion at 0 h. This is potentially due to quenching or interfering effects of the iron oxide on the fluorescent dye. When we undertook fluorescent analysis of cell uptake we did not observe any significant impact on the conclusions drawn.

### *In vitro* Magnetically Driven Cellular Uptake of Nanoemulsions

The cellular uptake of the DAMP4-PEG nanoemulsions and iron oxide loaded DAMP4-PEG nanoemulsions was evaluated using two different cell lines over time to assess whether a magnetic field could alter the uptake. The nanoemulsions were incubated with cells for 0.5, 2 or 4 h in the presence or absence of a magnetic field. RAW 264.7 cells, an immortalized murine macrophage cell line, were chosen to understand the immune evasion properties of the unloaded and iron oxide loaded DAMP4-PEG nanoemulsions. In the absence of a magnetic field the cell uptake was slow and there was no significant difference between unloaded or iron oxide loaded nanoemulsions (**Figure 5A**). This demonstrates that the PEGylation of DAMP4-PEG provides sufficient immune evasion to avoid clearance by macrophages of the reticuloendothelial system.

The RAW 264.7 cells demonstrated significantly increased uptake of iron oxide loaded DAMP4-PEG nanoemulsions at all time points in the presence of a magnetic field compared to iron oxide loaded DAMP4-PEG nanoemulsions in the absence of a magnetic field (**Figure 5A**). There was also a significant difference at all time points between the DAMP4-PEG nanoemulsions in the presence of a magnetic field. These two results indicate that the magnetic field ‘switched on’ the uptake of the iron oxide DAMP4-PEG nanoemulsion. We propose that the magnetic field induces the iron oxide nanoemulsion to move and rest on the cell membrane. This process removes the Brownian motion exhibited by nanoemulsions in solution that acts as a kinetic barrier to nanoparticle uptake. Thus, the nanoparticles are in constant close proximity of the cell membrane promoting continuous uptake dictated by the kinetics of endocytic pathway mechanisms. The enhanced uptake of the iron oxide nanoemulsion facilitated by magnetic field was capable of being sustained over time with the median fluorescent intensity increasing from 146970 to 454477 at 0.5 and 4 h respectively. This result shows that RAW 264.7 macrophages have significant capacity to continuously uptake and process nanoparticles. This is of clinical relevance with the revelation of a dosage threshold for Kupffer cell (a type of phagocytic liver macrophage) clearance of nanoparticles in the liver that once overcome may enhance the efficacy of nanomedicine.[21] We also observed using confocal microscopy many endosomal granular bodies within the cells and fluorescent signal within the RAW 264.7 cells treated with the iron oxide loaded DAMP4-PEG nanoemulsion in the presence of a magnetic field for 2 h (**Figure 5B**). To a lesser extent the granular bodies could be seen for RAW 264.7 cells treated with the iron oxide loaded DAMP4-PEG nanoemulsion when in the absence of a magnetic field for 2 h corresponding to the reduced uptake reported.

Next, the cell uptake of nanoemulsions by SKOV3 a human ovarian adenocarcinoma cell line was evaluated in the absence and presence of an external magnetic field. The SKOV3 cells demonstrated a different profile for their cell uptake kinetics of nanoemulsions compared to the RAW 264.7 cell line. The uptake of iron oxide loaded DAMP4-PEG nanoemulsion in the presence of a magnetic field was not significantly different at the 0.5 h time point compared to either of the controls (**Figure 6A**). This supports our hypothesis that the iron oxide loaded DAMP4-PEG nanoemulsions are not pulled directly into the cell but rest on the cell membrane to await endocytic uptake. At the 2 and 4 h time points the iron oxide loaded DAMP4-PEG nanoemulsion in the absence of a magnetic field was significantly different compared to the iron oxide loaded DAMP4-PEG nanoemulsion in the presence of a magnetic field with median fluorescent intensities of 1871 ± 148 and 76642 ± 6054 at 2 h and 7691 ± 1394 and 318348 ± 63872 at 4 h. There was also a significant difference at 2 and 4 h for the DAMP4-PEG nanoemulsion and the iron oxide loaded DAMP4-PEG nanoemulsion under the influence of a magnetic field. The SKOV3 demonstrates an enormous potential for continuous and rapid uptake of large quantities of nanoemulsions, once they have overcome the initial lag in uptake, with median fluorescent intensities similar to that observed in RAW 264.7 cells. This delay indicates that different uptake mechanisms are at play for the RAW 264.7 macrophages and the SKOV3 cells. As previously mentioned RAW 264.7 macrophages are professional phagocytic cells that efficiently utilize phagocytosis and macropinocytosis to take in large foreign objects. This would allow them to once in close proximity with the nanoemulsion efficiently uptake large quantities rapidly. Whereas, the uptake of nanoparticles in SKOV3 cells would most probably go through caveolin and clathrin independent endocytic pathways. It has been demonstrated in the literature for SKOV3 cells that nanoparticles without a targeting ligand are typically taken through these pathways and not clathrin mediated endocytosis.[22] Caveolin endocytic pathways have limits on the size of objects that can be endocytosed; nanoparticles with a diameter >100 nm, such as those used in this study, are too large. Using confocal microscopy we observed large numbers of endosomal granular bodies associated with fluorescent signal within the SKOV3 cells treated with the iron oxide DAMP4-PEG nanoemulsion in the presence of a magnetic field (**Figure 6B**). The granular bodies were not readily observed within the cells when in the absence of a magnetic field or for DAMP4-PEG nanoemulsion.

**Figure 6.**
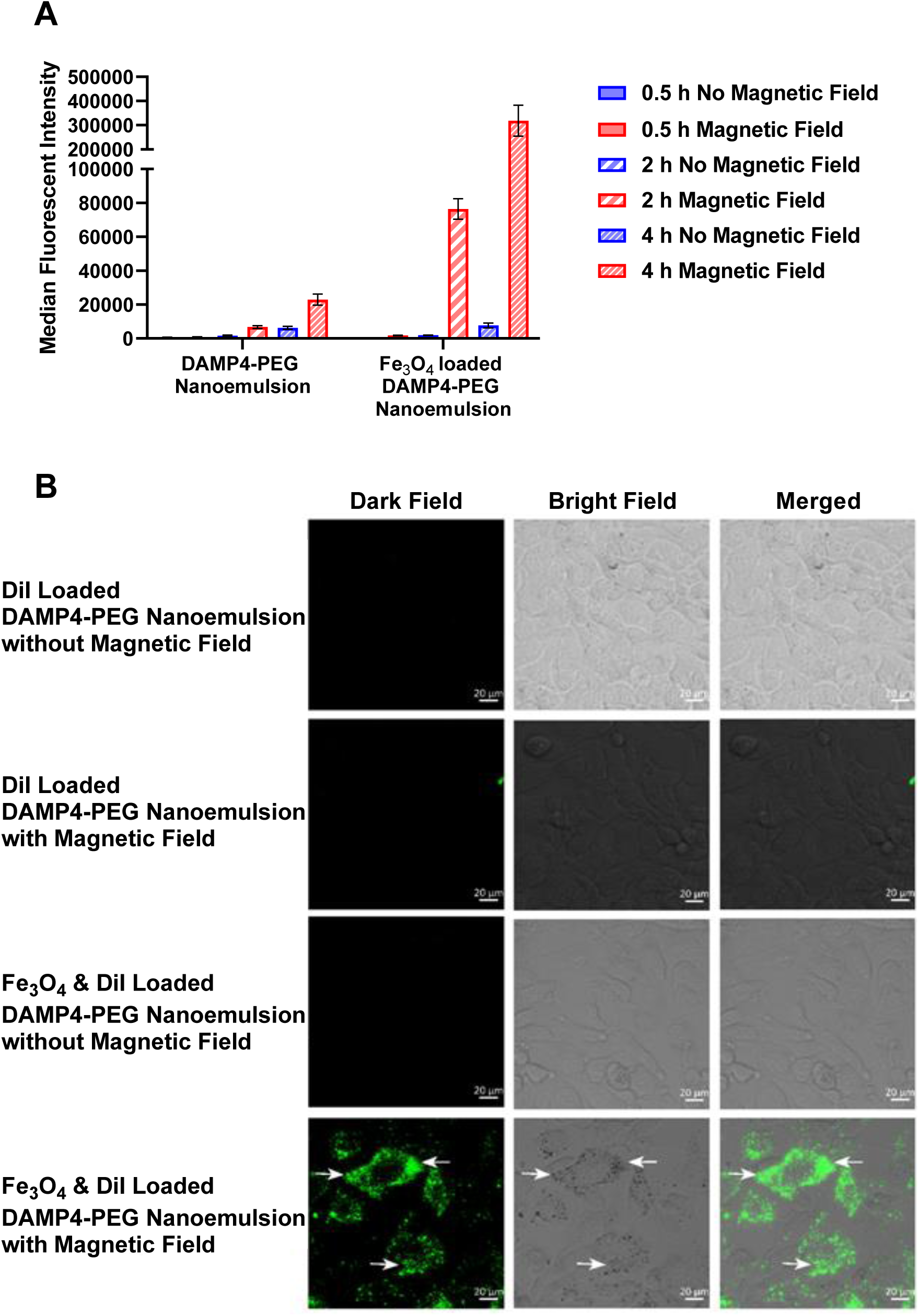
The cell uptake of DAMP4-PEG and iron oxide loaded DAMP4-PEG nanoemulsions by SKOV3 cells. **(A)** Flow cytometric data of the median fluorescent intensities at different time points in the presence and absence of a magnetic field. **(B)** Representative confocal images showing the cell uptake of DAMP4-PEG loaded nanoemulsions by SKOV3 cells in the presence and absence of a magnetic field after 2 h of incubation. Arrows indicate granular bodies. The mean + s.d. from 6 separate replicates is shown.

We evaluated the cell viability of three different cell lines after 0.5, 2 and 4 hours of incubation to investigate whether there was significant cell death over the course of the cell uptake experiments. DAMP4-PEG nanoemulsions and iron oxide loaded DAMP4-PEG nanoemulsions were incubated with three different cell types associated with the tumor microenvironment: RAW 264.7, SKOV3 and fibroblast (primary human mammary fibroblasts). The dosage of nanoemulsion added to the cells was kept constant. In addition, some cells were separately subjected to a magnetic field to promote uptake of the iron oxide loaded nanoemulsions. There was no significant difference in the cell viability for RAW 264.7 cells treated with either of the nanoemulsions in the absence of a magnetic field at all three time points (**Figure S3 A**). We observed a significant decrease in the viability of the RAW 264.7 cells treated with iron oxide loaded DAMP4-PEG nanoemulsion in the presence of a magnetic field compared to either the control and DAMP4-PEG nanoemulsion at all time points (**Figure S3 B**). This may be due to rapid magnetically driven uptake of iron oxide loaded nanoemulsions. Rapid uptake of iron oxide loaded DAMP4-PEG nanoemulsion would significantly increase the iron burden inside the cells with a subsequent spike in ROS potentially inducing cell death or limiting proliferation by widespread cellular damage. The cytotoxic effect of the iron oxide DAMP4-PEG nanoemulsion appeared to reduce over time as the cell viability increased from 54% to 76% at 0.5 and 4 h respectively. Heterogeneous uptake of iron oxide loaded nanoemulsions may result in some cells taking up significantly more iron oxide than others inducing apoptosis. As a result, cells that experience a reduced iron concentration from low iron oxide loaded nanoemulsion uptake or by sequestration of iron oxide in apoptotic bodies can continue to proliferate. The DAMP4-PEG nanoemulsions and iron oxide loaded DAMP4-PEG nanoemulsions in the absence of a magnetic field had no significant effect on the cell viability of SKOV3 cells compared to the control (**Figure S3 C)**. However, in the presence of a magnetic field the iron oxide loaded DAMP4-PEG nanoemulsion saw a significant decrease in the cell viability at 0.5 and 2 h, with a cell viability of 76% and 71% respectively (**Figure S3 D**). There was no significant difference between the controls and the iron oxide loaded DAMP4-PEG nanoemulsion at 4 h. This indicates some level of cytotoxicity derived from the uptake of iron oxide loaded DAMP4-PEG nanoemulsion by the cells during the assay. However, the magnitude of toxicity was less than that observed in the RAW 264.7 cells. The SKOV3 cells recovered at the 4 h time point with a normalized absorbance of 88% at 4 h at a similar level to the control and DAMP4-PEG nanoemulsion. This also indicates a more rapid recovering of cell proliferative ability than that seen in the RAW 264.7 cells. This difference in toxicity over time was probably due to the more rapid uptake by phagocytic uptake pathways observed in the RAW 264.7 cells compared to the SKOV3 cells. Finally, the cell viability of primary human mammary fibroblasts treated with DAMP4-PEG nanoemulsions and iron oxide loaded DAMP4-PEG nanoemulsions was not-significant in the absence of a magnetic field (**Figure S3 E**). In the presence of a magnetic field there was also no significant decrease in the viability observed at all 3 time points compared to the control (**Figure S3 F**). Fibroblasts are known to have extensive caveolae pits allowing for efficient caveolin assisted uptake, however, this process is size restricted to particles with a size less than 100 nm.[23] As discussed previously, both the DAMP4-PEG and iron oxide loaded DAMP4-PEG nanoemulsions have a hydrodynamic radius of approximately 175 and 250 nm when coated with serum proteins making them too large for efficient uptake by caveolin pathways. Fibroblast uptake of the nanoemulsion would then have to be regulated by either a non-professional phagocytic pathway, that is much less efficient than that seen in macrophages such as RAW 264.7, or caveolin and clathrin independent pathways.[24]

## Conclusion

We have reported the facile preparation of magnetic biosurfactant stabilized nanoemulsions dual encapsulating iron oxide nanoparticles and hydrophobic drug/dye for magnetic targeting of cancer cells. We systemically evaluated the key factors to stabilize iron oxide loaded nanoemulsions using four designed biosurfactants: AM1, SurSi, DAMP4 and DAMP4-PEG. The strong steric and electrostatic stabilization of DAMP4-PEG was imperative to ensure stability in biological media and overcome the interfacial behavior from loading iron oxide nanoparticles inside the nanoemulsion. Using the DAMP4-PEG platform, we showed that by using an external magnetic field that RAW 264.7 and SKOV3 cells could rapidly uptake DiI (fluorescent dye) and iron oxide loaded DAMP4-PEG nanoemulsions. Further, SKOV3 cells demonstrated a delayed uptake profile of magnetic nanoemulsions compared to RAW 264.7 cells. The cell uptake and cell viability studies demonstrate the advantages of using such multifunctional nanoemulsion drug delivery systems for magnetically targeted drug delivery. As nanoparticles arrive at the tumor site, the application of magnetic field could potentially increase uptake by cancer cells, thus increasing their cytotoxicity. This study offers a versatile nanoemulsion system that’s provides a foundation for the formulation of many unique nanoemulsion systems encapsulating different hydrophobic nanoparticles for drug delivery and diagnostic applications.

## Acknowledgements

The work was supported by the Australian Research Council Discovery Project (DP200101238). Zhao acknowledges the funding from the National Health and Medical Research Council Investigator Grant (APP2008698). Yang acknowledges the funding support from the Australian Research Council (Grant No. CE200100009). This work was performed in part at the Queensland node of the Australian National Fabrication Facility, a company established under the National Collaborative Research Infrastructure Strategy to provide nano- and micro-fabrication facilities for Australia’s researchers.

## Supplementary figures

**Table S1.**
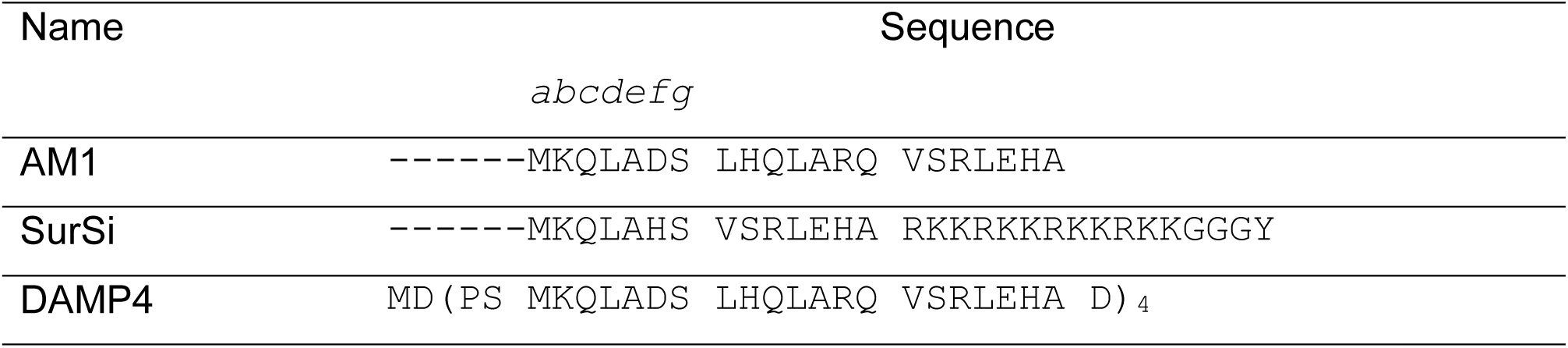
Alpha-helical peptide/protein sequences.

**Figure S1.**
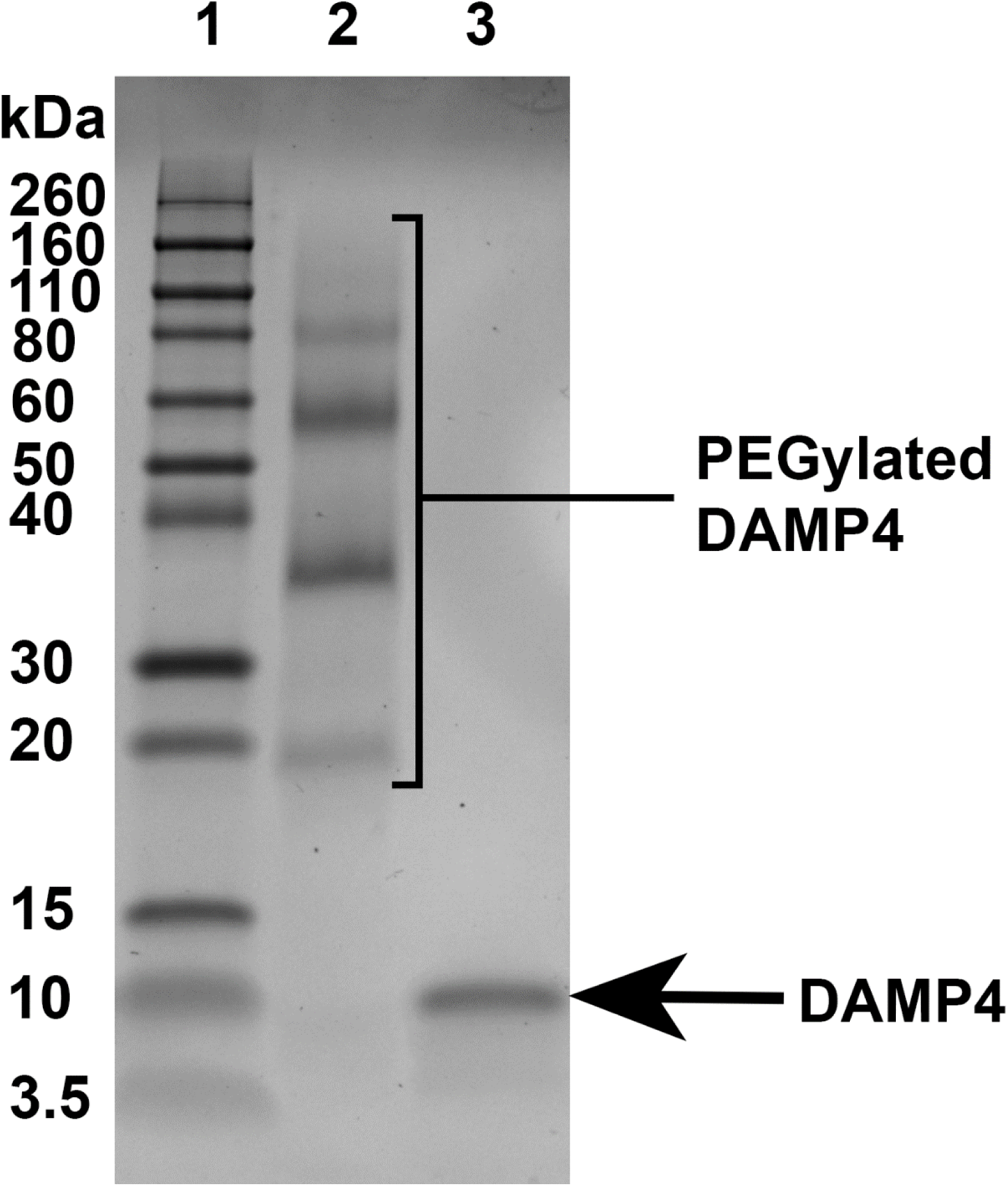
SDS-PAGE gel image showing the reaction between DAMP4 and 5 kDa NHS-PEG and DAMP4 before reaction. **Lane 1:** Novex™ Sharp Pre-stained Protein Standard, **Lane 2:** PEGylation reaction product, **Lane 3:** DAMP4 protein product.

**Figure S2.**
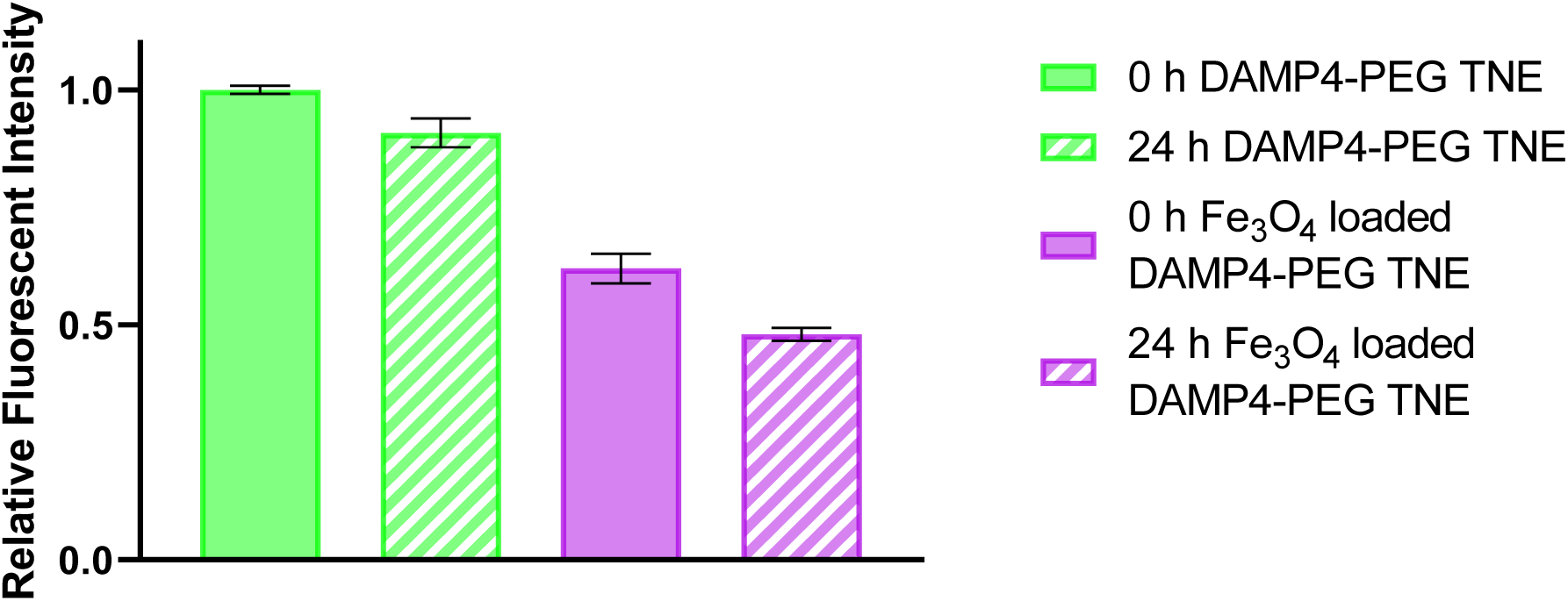
Dye release study of DiI loaded DAMP4-PEG stabilized nanoemulsions over 24 hours. The mean + s.d. from 3 separate replicates is shown.

**Figure S3.**
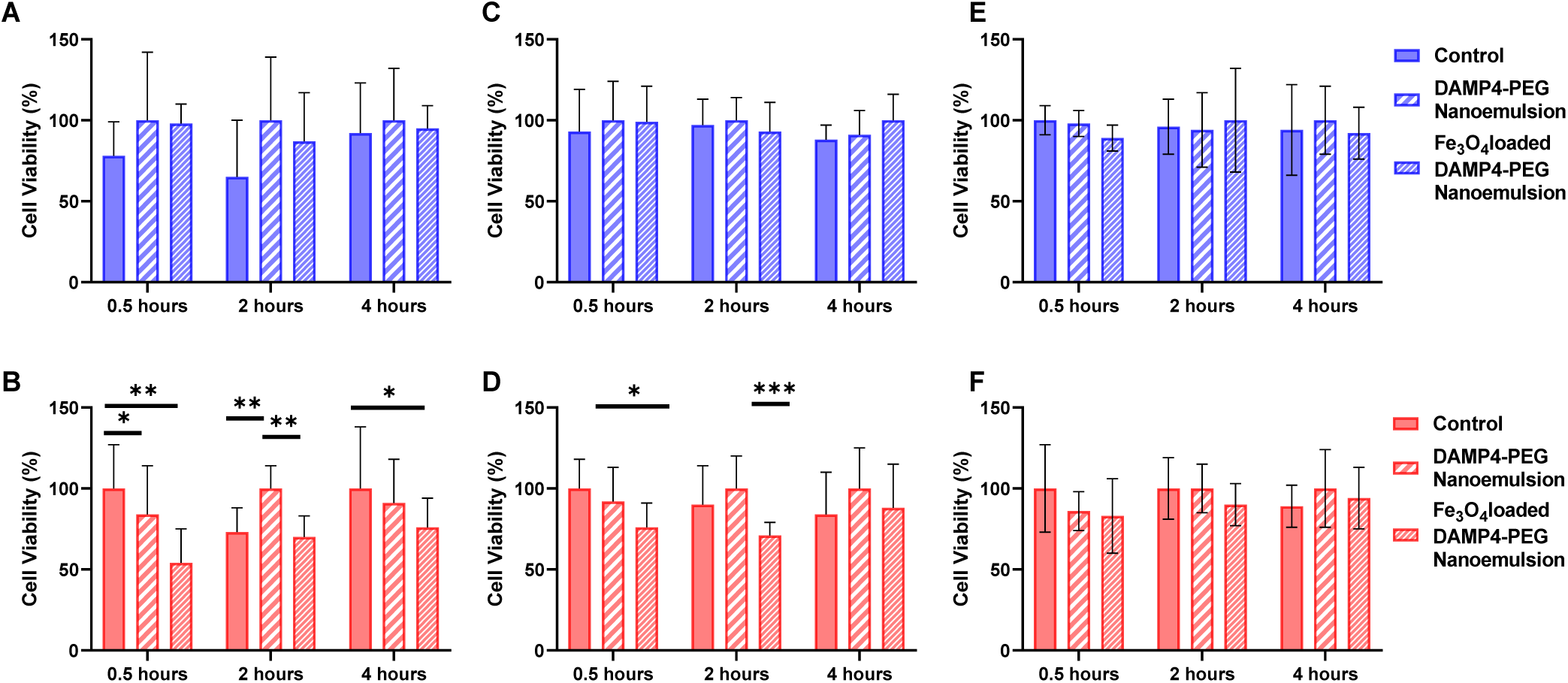
*In vitro* cytotoxicity of DAMP4-PEG nanoemulsions and iron oxide loaded DAMP4-PEG nanoemulsions. RAW 264.7 murine macrophage cells (**A**) in the absence of magnetic field and (**B**) in the presence of magnetic field. SKOV3 ovarian cancer cells (**C**) in the absence of magnetic field and (**D**) in the presence of magnetic field. Human mammary fibroblast cells (**E**) in the absence of magnetic field and (**F**) in the presence of magnetic field. The mean + s.d. from 10 separate replicates is shown.

## Notes

### Competing Interest Statement

The authors have declared no competing interest.

